# Improved Insights into Protein Thermal Stability: From the Molecular to the Structurome Scale

**DOI:** 10.1101/055897

**Authors:** Fabrizio Pucci, Marianne Rooman

## Abstract

Despite the intense efforts of the last decades to understand the thermal stability of proteins, the mechanisms responsible for its modulation still remain debated. In this investigation, we tackle this issue by showing how a multi-scale perspective can yield new insights. With the help of temperature-dependent statistical potentials, we analyzed some amino acid interactions at the molecular level, which are suggested to be relevant for the enhancement of thermal resistance. We then investigated the thermal stability at the protein level by quantifying its modification upon amino acid substitutions. Finally, a large scale analysis of protein stability - at the structurome level - contributed to the clarification of the relation between stability and natural evolution, thereby showing that the mutational profile of thermostable and mesostable proteins differ. Some final considerations on how the multi-scale approach could help unraveling the protein stability mechanisms are briefly discussed.

## 1. Introduction

To accomplish their biological function, the majority of proteins undergo a folding transition from a random coil state to a well defined three-dimensional folded structure. This process can be thermodynamically characterized by the standard folding free energy Δ*G*^0^, *e.g.* the difference in standard free energy between the folded and the unfolded states. If we assume that the pressure of the system and the other environmental variables (pH, ionic strength,…) are constant and that the change in heat capacity upon folding is temperature (*T*)-independent, and if we consider only two state reversible folding processes, the folding free energy can be written as:

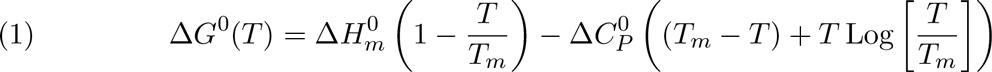

where *T_m_* is the melting temperature of the protein, and 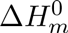 and 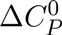 are the standard folding enthalpy at *T_m_* and the change in heat capacity upon folding, respectively. Throughout this investigation we will focus on **protein thermal stability** characterized by the descriptor *T_m_,* the temperature of heat denaturation defined as Δ*G*^0^(*T_m_*) = 0.

From a theoretical perspective, the investigation of the thermal stability properties of proteins is interesting to understand how organisms adapt to their environment. Living organisms can survive over a wide range of temperatures that go from below 0°C to above 100°C. Getting insight into the mechanisms used by proteins to modulate their thermoresistance is a way to better understand how extremophiles adapt to these extreme environments.

**Figure 1.**
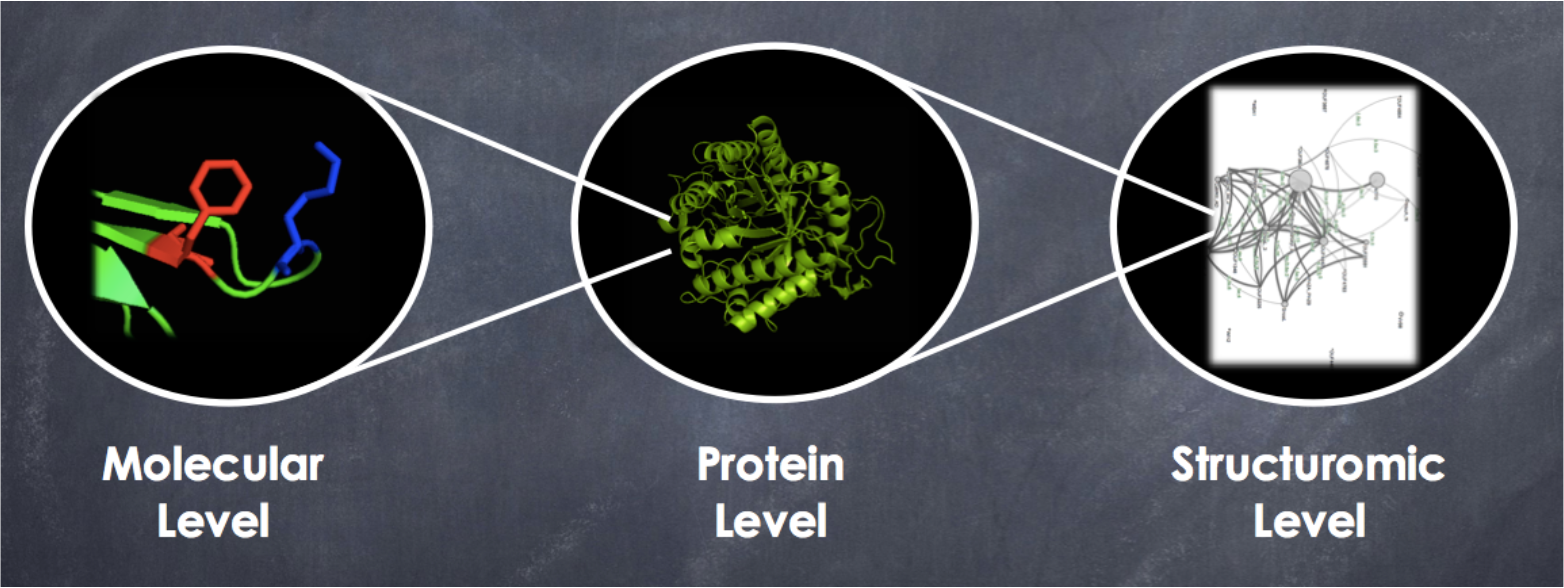
Schematic picture of the multi-scale nature of the protein thermal stability issue.

From a more applicative point of view, the optimization of biotechnological and bio-pharmaceutical processes often require proteins that work in conditions that are different from their physiological ones. Experimental protein design methods such as those based on direct evolution are only partially successful due to the vastness of the sequence space that needs to be scanned and to the low probability of obtaining thermally stabilizing mutations out of random mutations. It is thus mandatory to get more precise understanding of the problem and to design bioinformatics-based protein design methods with improved, faster and more trustable outcomes.

In this paper we tackled the protein thermal stability issue with a multi-scale approach, linking the molecular, macromolecular and structurome levels (Fig. 1). At the **molecular scale**, thermal stability is reflected in the *T*-dependence of the amino acid interactions which we analyzed through the development of *T*-dependent force fields. At the **protein scale**, we investigated and predicted the thermal stability of full protein structures, using the molecular-scale force fields. At the **large scale**, we considered the ensemble of proteins of known structure, the so-called structurome, and studied protein stability in an evolutionary context using the prediction methods designed at the protein level. In summary, the informations gathered at the molecular level were used at the protein level and conversely, and the protein-level information were in turn exploited at the structurome level.

## 2. Investigation method

It is of utmost importance in any multi-scale problem to choose an appropriate investigation method and simplification level. In particular, we cannot use a molecular-level approach that is too detailed, as it would make the analysis at the structurome scale too computationally expensive. Conversely, using methods derived from large-scale evolutionary information does not allow gathering information about the molecular scale.

An adequate level of simplification for studying multi-scale protein thermal stability involves using statistical mean-force potentials, derived from non-redundant sets of well-resolved protein structures [1, 2, 3, 4, 5]. The mean force potential Δ*W* of a sequence motif *s* (single amino acid or amino acid pair) adopting a conformational state *c* (inter-residue distance, backbone torsion angle domain, residue solvent accessibility, or their combination) is defined from the probabilities *P*(*c*, *s*), *P*(*c*) and *P*(*s*) through the inverse Boltzmann law:

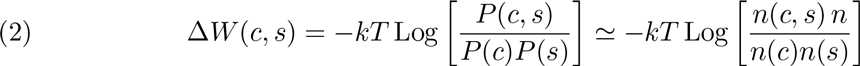

where *k* is the Boltzmann constant. These probabilities can be estimated in terms of the number of occurrences *n*(*c*,*s*), *n*(*c*) *n*(*s*) and *n* in a structure dataset (see [5] for details).

The potentials so defined conserve the memory of the protein dataset from which they are extracted. It is generally admitted that choosing a dataset that is large enough and well sampled, and that satisfies reasonable criteria in terms of structure resolution and pairwise sequence identity, yields well defined and informative potentials. It is however important to emphasize that the memory of statistical potentials is not only a disadvantage but also a unique strength: deriving potentials from proteins sharing a given characteristic, such as protein size [6] or thermostability [7], yields potentials that describe this characteristic. This advantage will be used in the next section.

Another advantage of these potentials is that they are fast enough to be applied on a large scale. They can be derived in a few minutes from standard datasets of thousands of protein structures. Their application to structure prediction or free energy computations is also quite rapid, which makes them an optimal tool to investigate the relation between stability and structure at the structurome scale. Note that these potentials are based on a simplified representation of the protein structure, in which only the main chain atoms and the average side chain centers are considered. In doing this we neglect the degrees of freedom of the side chains. This is of course a loss of information but it is the price that we have to pay if we want a methodology that can be applied on a large scale.

## 3. Analysis at the molecular level

The understanding of the molecular mechanisms that lead to the enhancement of protein thermoresistance is a longstanding problem in protein science [8, 9, 10, 11]. No unique or specific mechanism has been found to be the major driving force of thermal stabilization, which instead is reached by a complex balance of different factors. Only some general trends have been identified, that are frequently only observed inside protein families or are even specific to a protein.

For example, salt bridges and cation-*π* interactions that are close-range interactions between a positively charged residue, and a negatively charged or aromatic residue, respectively, seem generally to contribute to the enhancement of the thermoresistance. Instead, interactions between non-charged polar residues seem to disfavor it [12, 13, 14, 15]. Other structural factors that appear related to the thermal resistance are the protein rigidity and the higher packing efficiency [16, 17]. These structural features can be achieved for example by shortening some loops, increasing the number of atoms that are buried in the protein core, or filling some buried cavities to optimize the packing. Note that these are only general tendencies that do not always work.

The lack of knowledge about the temperature dependence of amino acid interactions makes the analysis of the thermal stability at the molecular level intricate. Usually, moreover, the studies of protein structure and stability are performed using force fields that do not take into account this *T*-dependence, which adds further uncertainty to the problem and the risk of misinterpreting the obtained results. We review here the technique that we used to get more insight into this dependence (for details we refer to [18, 7]) and apply it to the analysis of the anion-π interactions.

The idea developed in [18, 7] consists in taking advantage of the bias of the potentials towards the dataset from which they are extracted. We constructed a set of about 200 proteins with known melting temperature and 3-dimensional (3D) structure and divided it into different subsets according to the *T_m_*-values of their members. Then we derived statistical potentials from each subset as explained in the previous section. The potentials obtained in this way reflect in principle the thermal stability properties of the ensembles from which they are derived.

More precisely, we constructed three datasets: a mesostable set *S^M^* with proteins characterized by 35°*C* < *T_m_* ≤ 65°*C* and mean melting temperature 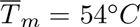, an intermediate set *S^I^* with 53°C < *T_m_* ≤ 73°*C* and 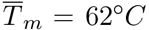, and a thermostable set *S^T^* with 65°*C* < *T_m_* ≤ 150°*C* and 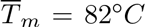. From each of them we derived the values of the statistical potentials at three different temperatures labeled as 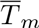, thereby extending Eq. (2):

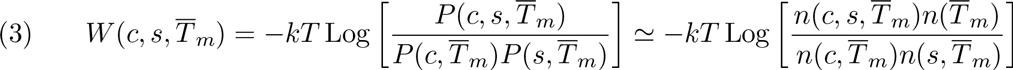

To illustrate de power of this approach, we analyzed anion-aromatic interactions between aspartic acid (Asp) and tyrosine (Tyr). Recent indications seem to point out that this kind of interactions could be important in the stabilization of protein structure since the positively charged edge of the aromatic residues could interact with the anion through an electrostatic anion-quadrupole interaction [29, 30, 31, 32].

Using our statistical potential methodology defined in Eqs (2, 3), with the sequence elements *s* being residue pairs and the conformational states *c* being interresidue distances between *C_µ_* pseudoatoms (defined as the average geometric centers of the heavy side chain atoms of a given residue type in the protein structure set), we derived the Asp-Tyr distance potentials. They are plotted in Fig. 2 as a function of the*C_µ_*-*C_µ_* distance at different values of the temperature (taken to be the 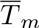 of the dataset from which they are derived), while in Fig. 3, a continuum extrapolation of the effect of the temperature on the anion-0 interaction is plotted.

We clearly observe in Figs 2–3 a stabilization effect due the temperature: the potential at higher temperature is shifted downwards with respect to that at lower temperature. Note that this does not mean that the Asp-Tyr interaction is more stable at higher *T*, but rather that it is more stable compared to the other residue-residue interactions. For all three temperatures, this interaction appears as destabilizing in the distance window between about 3 and 6 Å, even if at higher temperature (red curve) the destabilization effect is reduced with respect to the low temperature regime (blue curve). These destabilizing conformations have frequently an anion-*π* stacking geometry similar to the one plotted in the picture and also known as *η*6 geometry [29]. Despite their intrinsic destabilizing nature, their presence could be explained by an energetic compensation from other interactions that involve other residues in the polypeptide chain [30]. Indeed, the authors of [30] suggest that more complex geometries can bring stabilization, such as anion-*π*-cation or anion-*π*-*π* stacking geometries.

Instead, at about 6-7 Å, there is an energy minimum that corresponds to the anion-*π* hydrogen bond conformations similar to the one depicted] in Fig. 2, in which a hydrogen bond is established between the oxygens of the carboxylic group of the aspartic acid and the hydroxyl group of the tyrosine. The effect of the temperature is quite substantial at the minimum and makes it deeper of about −0.1 kcal/mol at higher *T*.

## 4. Analysis at the protein level

The informations about the amino acid interactions that we have derived using the *T*-dependent and standard statistical potentials are utilized in this section to better understand the stability properties at the protein level. More precisely, we are interested in finding the relations between the structural and energetic characteristics of the proteins and the change in their melting temperature *T_m_* upon amino acid substitutions:

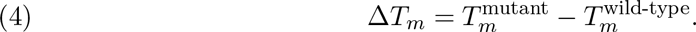

To evaluate quantitatively and automatically how the *T_m_* of a given protein changes upon point mutations, we developed a predictor called HoTMuSiC [27], which is freely available for academic use at dezyme.com.

There are two versions of the tool. In the first, only the protein structure is required as input. It uses 9 standard statistical potential terms 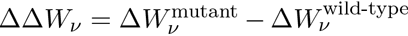, which differ in the sequence motifs *s* and conformational states *c* on which they are based (see Eq. (2)), with in addition an independent term and two volume terms Δ*V*± that describe the influence of the creation of a hole or a stress in the protein structure upon replacing a large into a small amino acid or conversely. These terms are summed with weight factors *α_i_*(***A***) chosen to be sigmoid functions of the solvent accessibility ***A*** of the mutated residue. The HoTMuSiC functional thus reads as:

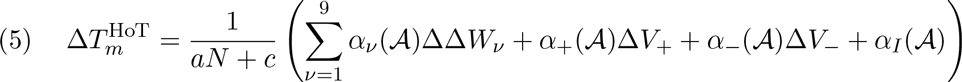

where *a*, *c* are two additional parameters and *N* is the total number of residues in the target protein. All the parameters are optimized using an artificial neural network so as to minimize the root mean square deviation between the predicted and the experimental values of Δ*T_m_* for a dataset (called here *MutS*) of about 1600 mutations introduced in 90 proteins, described in [25].

**Figure 2.**
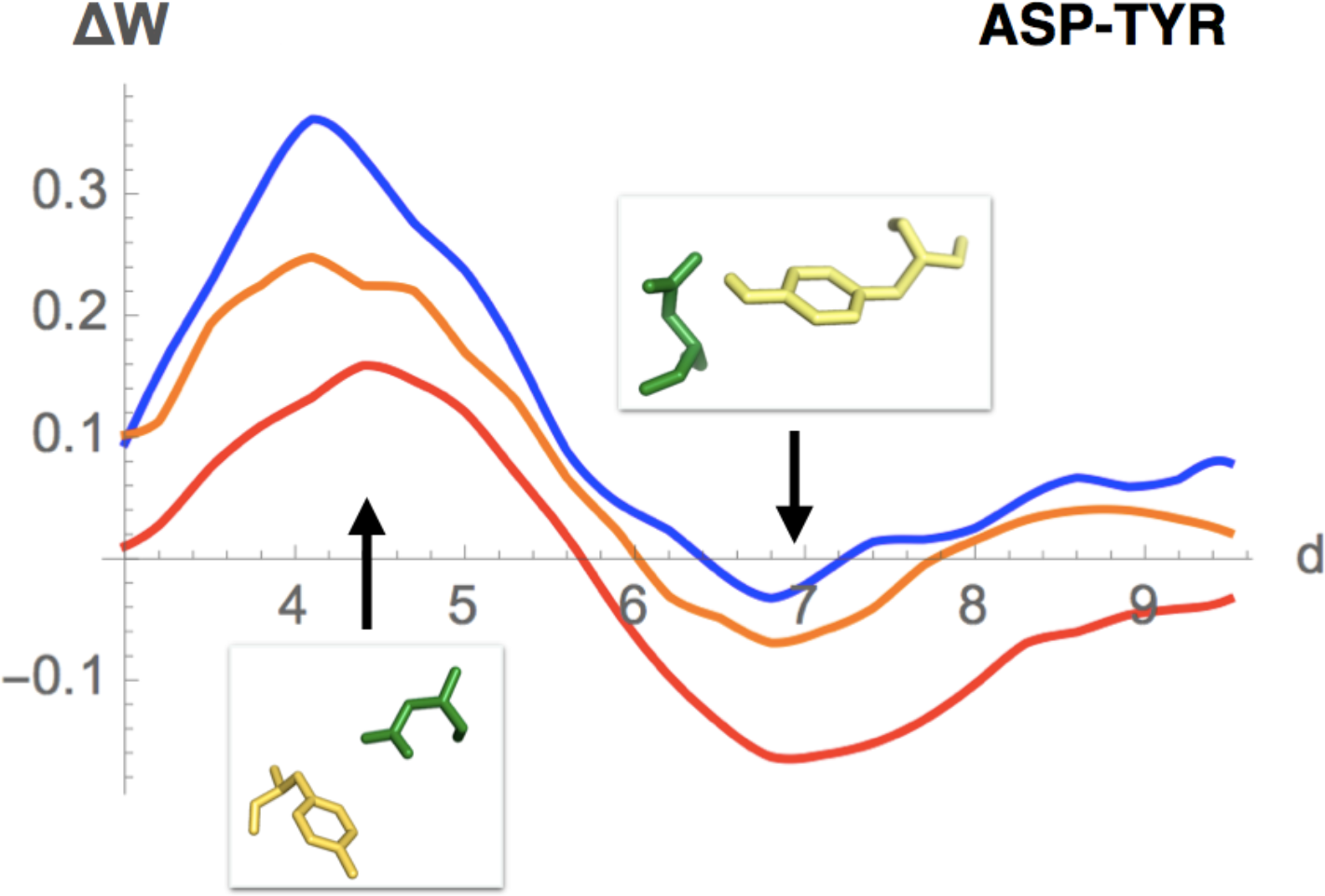
Asp-Tyr inter-*C_µ_* distance potential at different temperatures. The distance is in Å and the energy in kcal/mol. The mesostable potential is in blue 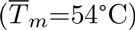, the thermostable one in red 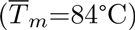 and the intermediate potential in orange 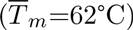. Typical geometries corresponding to the maximum (Asp 198 and Tyr 156 in 1IS9[28], inter-*C_µ_* distance of 4.8 Å) and to the minimum (Asp 155 and Tyr 142 in 1AKY [28], inter-*C_µ_* distance of 7.1 Å) are depicted.

In the second HoTMuSiC version, called *T_m_*-HoTMuSiC, the informations derived about the thermal stability at the molecular level were taken into account. The *T_m_* of the wild-type protein is here required in addition to its structure, and the Δ*T_m_* is computed by adding *T*-dependent statistical potential terms defined in Eq. (3) to the functional of Eq.(5):

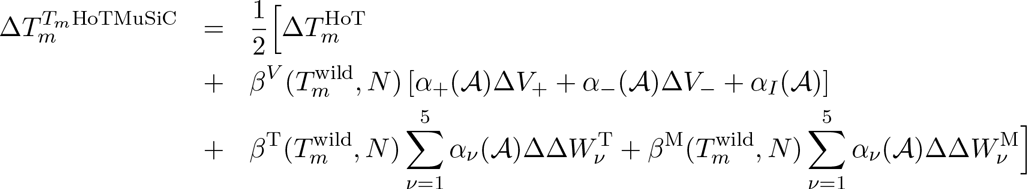

**Figure 3.**
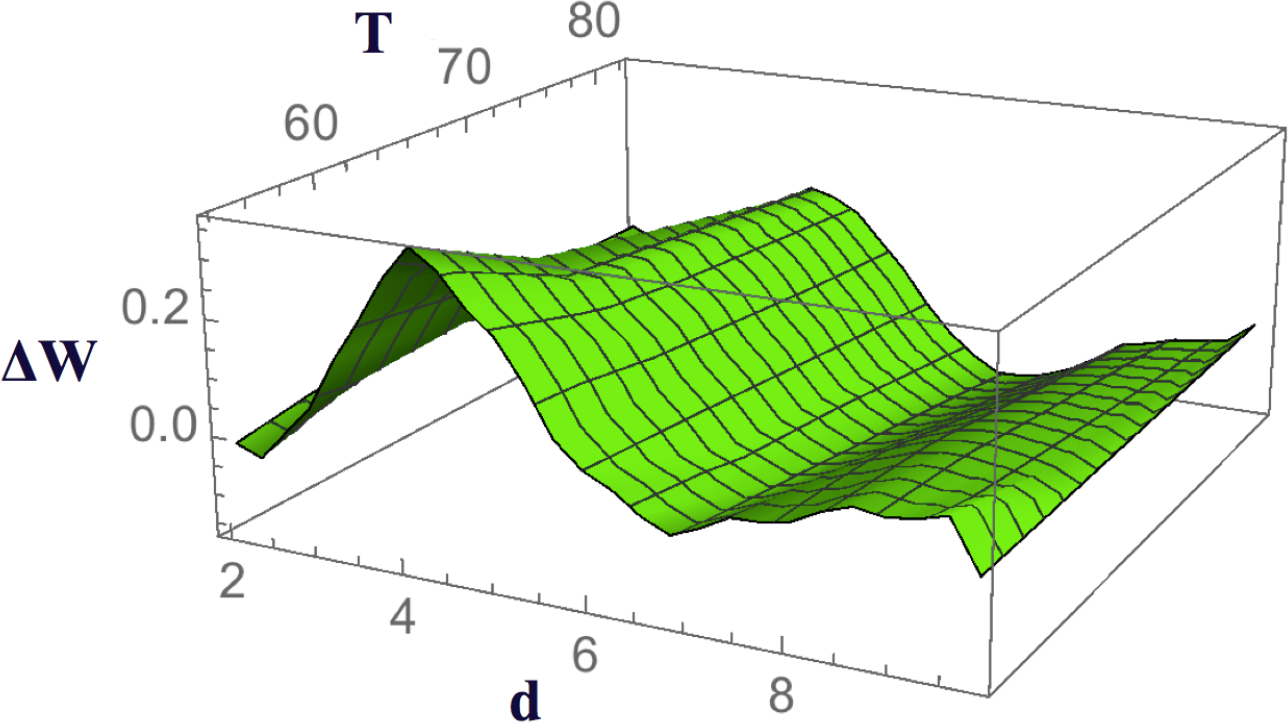
Asp-Tyr potential (in kcal/mol) as a function of the inter-*C_µ_* distance (in A) and the temperature (in °C), obtained from a continuum extrapolation of Fig. 2.

The 5 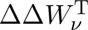 and 5 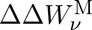 terms are folding free energy changes computed from the sets *S^T^* and *S^M^* of thermostable and mesostable proteins (defined in section 3), respectively, with various values of *s* and *c*; *β^T^*, *β^M^* and *β^V^* are parabolic functions of the melting temperature and the number of residues of the target protein (see [27] for details).

The scores of HoTMuSiC and *T_m_*-HoTMuSiC were evaluated in strict 5-fold cross validation on the above cited learning set *MutS* of about 1600 mutations. They are quite good: 4.3°C for HoTMuSiC and 4.2°C for *T_m_*-HoTMuSiC, which decreases to 2.9°C for both methods when 10% outliers are removed. These performances are much better than those of other methods with similar approximation levels [27]. As expected, the predictions from *T_m_*-HoTMuSiC are more accurate than those from the standard HoTMuSiC, given that additional potentials are considered, which take more properly into account the thermal stability properties of the proteins. However, the difference in performance is smaller than expected, especially in the light of the results shown in the next section where a relation between the *T_m_* of the wild type protein and the Δ*T_m_* distribution of point mutations is observed. This can be attributed to the fact that the *T*-dependent potentials are extracted from small datasets of about hundred proteins and thus are noisy, even if some tricks were employed to limit this small-size effect [27].

## 5. Analysis at the structuromic scale

On the basis of the results obtained at the protein level, we analyzed the thermal properties of proteins at the structurome level in view of understanding how the physical principles that drive the thermal optimization are reflected in the evolutionary pressure on the sequences, which occur in response to the thermal conditions of the environment. In the last decade a series of investigations tackled this issue (see for example [19, 20, 21, 22, 23, 24] and reference therein), but the answers remained elusive and too frequently model dependent. Some crucial questions about the relation between thermal stability and evolution remain open:

- Are the effects of amino acid mutations on the protein stability conserved during natural evolution? Are they universally distributed ?
- How is the site-specific evolutionary rate linked to the structural characteristics of the proteins and to their thermal stability properties ? What is the relation between these properties and the functional constraints ?

We focused on the first point. It is still debated whether or not the distribution of the effects of amino acid mutations on protein stability is universal and whether or not they are conserved during evolution. Some investigations seem to indicate that they are, while others suggest that the distribution is universal only across proteins belonging to bacterial and eukaryote organisms, with some (unexplained) deviation in the archeon [19, 20, 21].

We analyzed the distribution profile of the experimental Δ*T_m_* values of the *MutS* dataset of about 1600 experimentally characterized mutations, as a function of the thermal stability properties of the wild type proteins. For that purpose, we clustered all the entries of the *MutS* dataset into four groups, as shown in Table 1. We clearly observe that the more thermostable the protein, the easier to destabilize with point mutations. Indeed, the mean 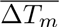 for the point mutations introduced in thermostable proteins 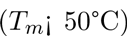 is 3°C smaller compared to those of hyperthermostable proteins 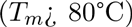. This effect is increased to about 4°C for the mutations introduced in the protein core due to the fact that thermostable proteins have usually a more compact structure. Instead, in the surface region, the difference between thermostable and mesostable proteins is reduced, even though the 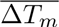 remains significant: about 1.5°C.

We used the protein-level predictor HoTMuSiC introduced in section 4 to evaluate the changes in stability of these 1600 mutations. As shown in Table 1, the mean 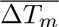 are almost identical to the experimental ones. Only the root mean square deviations are smaller than the observed ones.

Motivated by these results, we started a large scale analysis employing HoTMuSiC in order to find other indications about the non-universality of the distribution. For that purpose, we chose with the help of PISCES [26] a set of about 25000 proteins derived from the whole PDB Data Bank [28], imposing a threshold of 95 % on the pairwise sequence identity and considering only X-ray protein structures with resolution ≤2.5 Å. We evaluated the Δ*T_m_* for all possible point mutations - i.e. 19 mutations per residue - for all proteins that belong to this dataset and that are hosted by one of the four mesophilic and four thermophilic organisms cited in Table 2, and analyzed the Δ*T_m_* distribution. We found that the mean 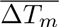 value for all the mutations introduced in mesophilic proteins is about −4.86°C and has to be compared with the analogous quantities for the proteins in the termophilic set that is −5.31 °C. In Fig. 4 the two predicted distributions are drawn. They are significantly different as measured by a P-value < 10^−10^.

The first thing to note about these results is that the destabilization effect is in general larger compared to the experimental values reported in Table 1. This is due to the fact that the experimentally characterized mutations are not random but are usually designed to improve the thermal stability of the protein. Instead, the systematic introduction of all possible mutations that can occur in a protein yields more destabilizing mutations on the average.

The second point is the small difference between the mutations introduced in proteins belonging to mesophilic and thermophilic organisms. This is probably due to the fact that, while thermophilic organisms host only thermostable proteins, mesophilic organisms host both mesostable and thermostable proteins.

**Table 1.** Mean values 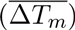 and root mean square deviations (σ) of Δ*T_m_* (in ^°^*C*) for the mutations inserted in proteins with different thermal characteristics. The experimental values are derived from the original literature and are reported in a dataset [25]. The computed values are obtained using *T_m_*-HoTMuSiC [27].

| Thermal stability<br>wild type | Solvent<br>accessibility | $\overline{\Delta T_m}$<br>exp | $\sigma$<br>exp | $\overline{\Delta T_m}$<br>pred | $\sigma$<br>pred |
| --- | --- | --- | --- | --- | --- |
| $T_m < 50^\circ\text{C}$ | | -1.4 | 4.6 | -1.3 | 2.3 |
| $50^\circ\text{C} < T_m < 65^\circ\text{C}$ | | -2.2 | 4.6 | -2.6 | 3.1 |
| $65^\circ\text{C} < T_m < 80^\circ\text{C}$ | | -3.2 | 4.7 | -2.8 | 2.9 |
| $T_m > 80^\circ\text{C}$ | | -4.4 | 5.6 | -4.3 | 3.4 |
| $T_m < 50^\circ\text{C}$ | Core | -1.5 | 5.1 | -1.4 | 2.6 |
| $50^\circ\text{C} < T_m < 65^\circ\text{C}$ | Core | -4.1 | 6.3 | -4.2 | 3.4 |
| $65^\circ\text{C} < T_m < 80^\circ\text{C}$ | Core | -4.6 | 5.2 | -4.3 | 3.1 |
| $T_m > 80^\circ\text{C}$ | Core | -5.4 | 6.2 | -5.7 | 3.3 |
| $T_m < 50^\circ\text{C}$ | Surface | -1.3 | 3.4 | -0.9 | 1.4 |
| $50^\circ\text{C} < T_m < 65^\circ\text{C}$ | Surface | -0.7 | 4.2 | -1.2 | 1.9 |
| $65^\circ\text{C} < T_m < 80^\circ\text{C}$ | Surface | -2.0 | 3.9 | -1.4 | 1.9 |
| $T_m > 80^\circ\text{C}$ | Surface | -2.9 | 4.0 | -1.9 | 2.0 |

**Table 2.** Mean values 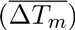 and root mean square deviations (σ) of Δ*T_m_* (in °C) for all possible mutations inserted in proteins of well-resolved X-ray structure and limited pairwise sequence identity, and belonging to the mentioned organisms. They are predicted using HoTMuSiC [27]. The last column contains the number of tested mutations.

| Host Organism | $T_{env}$ | Domain | $\overline{\Delta T_m}$ | $\sigma$ | $N_{mut}$ |
| --- | --- | --- | --- | --- | --- |
| Pyrococcus Furios | 95°C | Archea | -5.22 | 5.03 | $1.84 \times 10^5$ |
| Methanococcus jannaschii | 85°C | Archea | -5.25 | 5.03 | $1.75 \times 10^5$ |
| Sulfolobus solfataricus | 80°C | Archea | -5.17 | 5.07 | $2.37 \times 10^5$ |
| Thermus thermophilus | 70°C | Bacteria | -5.34 | 5.02 | $14.7 \times 10^5$ |
| Bos taurus | 39°C | Eukaryote | -4.94 | 4.92 | $11.7 \times 10^5$ |
| Homo sapiens | 37°C | Eukaryote | -4.80 | 4.95 | $178.2 \times 10^5$ |
| Mouse | 37°C | Eukaryote | -4.79 | 5.12 | $58.0 \times 10^5$ |
| E. Coli | 37°C | Bacteria | -5.06 | 4.95 | $27.4 \times 10^5$ |

**Figure 4.**
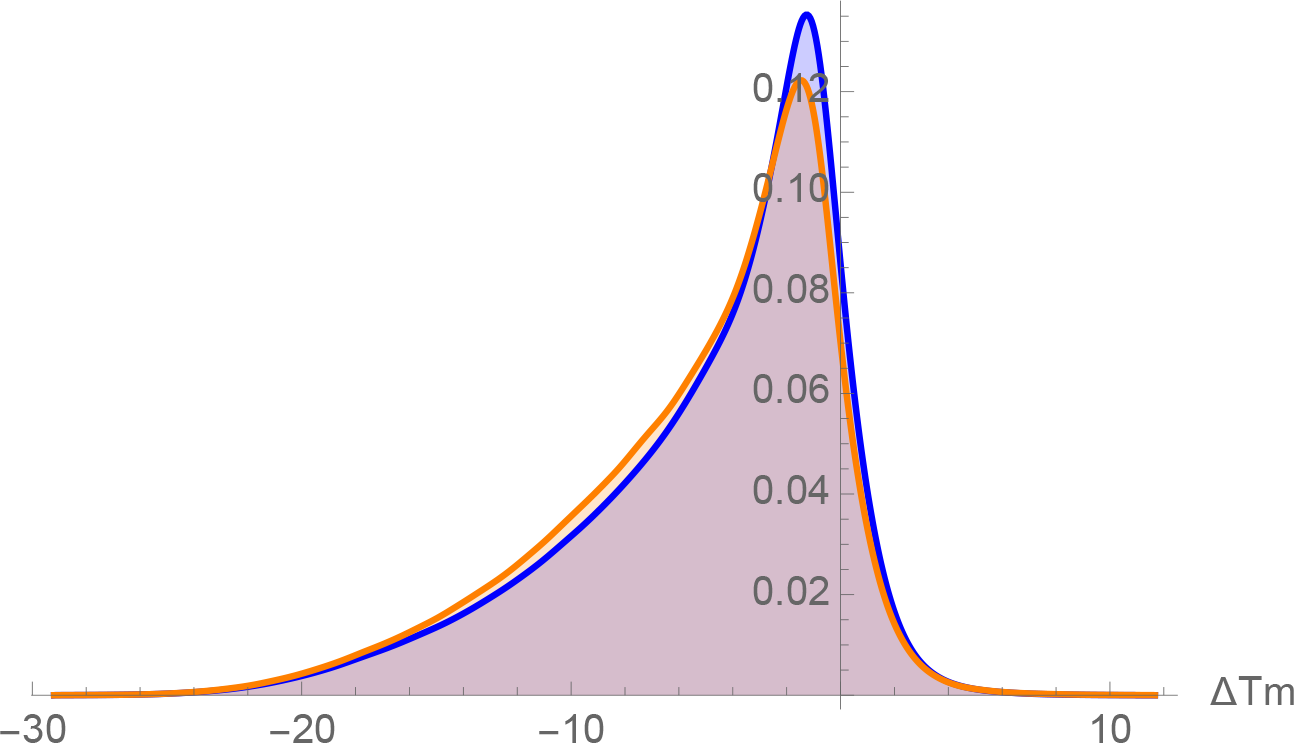
Predicted Δ*T_m_* distributional profile (in °C) for all possible mutations inserted in proteins belonging to mesophilic organisms (blue) or in thermophilic organisms (orange).

## 6. Conclusion

In this paper we have shown how a multi-scale approach can be utilized to study protein thermal stability. The methods employed in this analysis, namely the statistical potentials and their temperature-dependent generalization, allow the exploration of different scales since they were chosen to satisfy a tradeoff between the accuracy at the molecular level and the fastness (and thus the applicability) on a large, structuromic, scale. In summary:

- **Molecular scale**. Extending some previous works, we have shown that some interactions can contribute differently to protein stability in the high and low temperature regimes. We focused on the anion-*π* Asp-Tyr interactions and found an overall (relative) stabilization effect of the temperature, especially for particular kinds of geometries.
- **Protein scale**. We briefly reviewed the HoTMuSiC construction, a tool to predict the change in thermal stability properties upon amino acid substitutions, in order to show how the molecular level information can be employed to study protein thermal characteristics.
- **Structuromic scale**. Utilizing HoTMuSiC we have shown that the effects of mutations on the protein stability seem to be related to the thermal characteristics of the wild type protein.

In the “omic” era, the multiscale approaches that become possible by the large amount of available experimental data can shed new light on the protein stability issue and allow a better comprehension of a wide series of phenomena. We will continue our efforts in this direction since this issue deserves further investigations. In particular, it would be interesting to establish closer connections between the different scales. More precisely, the use of the thermal stability properties at small scale encoded in the temperature-dependent statistical potentials as well as all the evolutionary and functional information obtained at large scale could be better employed to improve the predictions of the change in *T_m_* upon mutations. Furthermore, the fast methods developed at the protein level could be fruitfully applied at large scale in order to improve the evolutionary models and the understanding of the intricate relation between stability, evolution and functional constraints in the protein universe.

